# TP53 knockout in MCF7 breast cancer cells induces sensitivity to fluoropyrimidine drugs

**DOI:** 10.1101/2022.02.10.479997

**Authors:** JM Warrington, S Mehta, VF Moy, JR Patel, C Liu, AR Rago, C McTighe, I Delgado, CE Banister, WH Gmeiner, PJ Buckhaults

## Abstract

TP53 mutations are present in all molecular subtypes of breast cancer and often correlate with decreased survival; however, few therapeutic options exist for patients with TP53-mutant breast cancers. To discover therapeutic strategies for these patients, we investigated the sensitivity of 129 FDA-approved chemotherapies to TP53-KO and TP53-WT MCF7 breast adenocarcinoma cells and found p53 loss to confer sensitivity to 5-fluorouracil (5-FU). We then treated the p53-null cells and isogenic controls with F10, a second-generation polymeric fluoropyrimidine, and found this preferential cytotoxicity of TP53-KO cells to be significantly magnified. F10 killing could only minimally be rescued by addition of exogenous uridine, whereas it was completely abrogated by addition of exogenous thymidine, suggesting DNA incorporation to be central to the cytotoxic mechanism of action. Furthermore, F10 killing of p53-null cells was persistent even in heterogeneous cellular mixtures more reflective of natural tumor evolution. Together, our results suggest F10 may widen the therapeutic window for TP53-mutant breast cancer by enhancing genotoxicity in cells resistant to apoptosis.

## Background

Breast cancer is the second most common cancer among women and is characterized by a high degree of histological and genetic heterogeneity (Bray et al., 2018; Polyak, 2011). Breast cancers can be grouped into several molecular subtypes according to the PAM50 classification scheme, which stratifies tumors based on the presence of biological markers such as hormonal receptors (ER and PR), human epidermal growth factor receptor 2 (HER2), and Ki67 (Parker et al., 2009). The molecular subtypes of breast cancer inform choices between therapeutic regimens; however, the clinical outcomes and toxicity profiles still vary drastically between patients with similar disease and treated with the same medication (Waks & Winer, 2019; van’t Veer et al., 2002; Tecza et al., 2018). There is a critical need for the development of new therapies with outcomes linked to specific genetic changes seen in breast cancers.

One such genetic aberration observed in breast cancers is TP53 mutation. TP53 is the most commonly mutated gene in human cancers, and The Cancer Genome Atlas (TCGA) Pan-Cancer Cohort documented TP53 loss of function in nearly 30% of all breast cancers via inactivating somatic mutations in both alleles (Kandoth et al., 2013; Donehower et al., 2019). The wild-type p53 protein causes cell cycle arrest and/or apoptosis in response to DNA-damage, hypoxia, or oncogene activation and its inactivation leads to unchecked cellular proliferation, even in the presence of DNA-damaging chemotherapies (Vousden & Lane, 2007; Ferreira et al., 1999).

TP53 mutations are unevenly distributed among the molecular subtypes of breast cancer. The gene is mutated in 80% of triple-negative breast cancers and 70% of HER2-enriched cancers but much less frequently in luminal B (30%) or luminal A (10%) cancers (Dumay et al., 2013). TP53 mutations may be late events (following PTEN loss) in basal-like tumor types, whereas they may be early gatekeeping events contributing to chemotherapy resistance in luminal A or B tumors (Martins et al., 2012; Silwal-Pandit et al., 2017). Despite this clear context-dependent role of TP53 mutation in breast cancer, survival outcomes indicate similarity in the clinical impact of TP53 mutation in several subtypes. In particular, TP53 mutations in luminal B, HER2-enriched, and normal-like tumors were found to be significantly associated with increased mortality in these patients, whereas weaker correlations between TP53 mutation and outcome are observed for basal-like tumors (Silwal-Pandit et al., 2014). TP53 mutations clearly provide evolutionary benefit to many tumor types and contribute to poor clinical outcome, so novel therapeutics exploiting these same TP53 mutations as an Achilles heel could significantly benefit patients with certain subtypes of breast cancer.

To directly characterize the impact TP53 mutations have on cancer therapeutics, without the confounding variables inherent to correlation studies, we generated TP53-knockout derivatives of MCF7 breast cancer cells and screened 129 FDA-approved chemotherapy drugs (NCI Approved Oncology Drug Set IV) against wildtype (TP53-WT) and knockout (TP53-KO) cells and identified drug responses in vitro that were caused by the introduced TP53 mutations. While some drugs were less effective on p53-null cells, we observed that several antimetabolites preferentially killed TP53-mutant cells. In particular, 5-fluorouracil (5-FU) showed the most significant TP53 mutation-associated increase in sensitivity in the AOD set IV. We next examined the TP53-dependent sensitivity of a novel polymeric fluoropyrimidine, F10, and observed an even more dramatic increase in sensitivity in p53-null cells relative to p53-WT MCF7s. This TP53-KO induced sensitivity to F10 could not be blocked by exogenously supplied uridine but was completely abrogated by exogenous thymidine, indicating the primary mechanism of action of F10 to be incorporation into DNA and likely generation of catastrophic DNA damage. These results suggest that unlike 5-FU, F10 may provide an increased therapeutic window for TP53-mutant breast cancers by limiting RNA-dependent toxicity and enhancing DNA-dependent genetic damage in cells unable to undergo cell cycle arrest.

## Methods

### Cell line culture

All human MCF7 breast adenocarcinoma (ATCC HTB-22) and HEK293T (ATCC CRL-3216) cell populations were maintained at 37°C, 5% CO2 in DMEM (Gibco, Cat. No. 11995-065) with 100ug/mL penicillin & 100ug/mL streptomycin (Sigma, Cat. No. P4333), and 10% FBS. MCF7 cells were additionally supplemented with 50mM Sodium pyruvate (Sigma, Cat. No. S8636), 1% GlutaMAX (ThermoFisher, Cat. No. 35050061), and 10ug/mL insulin. All cells were passaged every 4 to 7 days to maintain sub-confluence and maintained in culture for a maximum of 30 passages.

### Generation of pooled and clonal TP53-KO MCF7 cell lines via CRISPR-Cas9

Pooled TP53-KO MCF7 cells were generated as previously published (Liu et al., 2018). In brief, human codon-optimized Streptococcus pyogenes wild-type Cas9 (Cas9-2A-GFP) was obtained from Addgene (Cat. No. 44719). Two chimeric guide RNA expression cassettes containing two of the following sgRNAs (sgRNA1: 5’-CCATTGTTCAATATCGTCCG-3’, sgRNA2: 5’-GACGGAAACCGTAGCTGCCC-3’, sgRNA3: 5’-TGGTTATAGGATTCAACCGG-3’) were ordered as gBlocks from IDT. gBlock1 contained sgRNA1 & sgRNA2 and gBlock2 contained sgRNA1 & sgRNA3. These gBlocks were amplified by PCR using the following primers: gBlock_Amplifying_F: 5’-GTACAAAAAAGCAGGCTTTAAAGG-3’ and gBlock_Amplifying_R: 5’-TAATGCCAAACTTTGTACAAGAAAGC-3’, after which PCR products were purified using Agencourt Ampure XP PCR Purification Beads obtained from Beckman Coulter (Cat. No. A63882).

1μg of Cas9 plasmid and 0.3μg of either gBlock were then cotransfected into MCF7 cells with Lipofectamine3000 (Invitrogen, Cat. No. L3000001) in a 6-well plate. Knockout cells created using gBlock1 were named Pool KO5.6 and knockout cells created using gBlock2 were named Pool KO3.4. After transfection, pure TP53-KO populations were selected by culturing cells in 10μM nutlin-3a (SelleckChem, Cat. No. S8059). Media containing nutlin-3a was changed every 3 days and cells were passaged every 6 to 8 days. After 2 months, clonal cell lineages were isolated from TP53-KO and TP53-WT control mixtures via limiting dilution in a 96-well plate. These cells were expanded for 2 weeks to obtain TP53-KO Clone 5.6, TP53-KO Clone 3.4, and a TP53-WT control clone.

### Drug screening in pooled TP53-KO MCF7 cells

MCF7 TP53-WT and TP53-KO Pools 5.6 and 3.4 were plated at a density of 5000 cells/well in polystyrene, flat-bottom 96-well plates. All 129 compounds from the NCI Approved Oncology Drugs Set IV (AOD-IV) were dissolved in DMF or DMSO at 10mM stocks. F10 was dissolved in PBS at a 300uM stock (Liao et al., 2005). TP53-KO and TP53-WT MCF7 cells were treated with all drugs from the 129-member set at concentrations ranging from 156.25nM to 10μM. F10 concentrations ranged from 72.34pM to 300uM. After 10 days, resazurin (Sigma, Cat. No. R7017) was added to each well and incubated at 37°C in a 5% CO2 incubator for 4 to 6 hours. A microplate reader then took optical measurements (ex: 535nm / em: 585nm) to quantify cellular viability. Area under the curve (AUC) from dose response curves were calculated with the auc function from R package “flux” to represent cell viability and hierarchical clustering and heatmap generation were performed using the heatmap.2 function in R package “gplots.” The volcano plot was produced with the R package “EnhancedVolcano.”

### Thymidine and Uridine Rescue Experiments

MCF7 TP53-WT and TP53-KO Clones 5.6 and 3.4 were plated at a density of 5000 cells/well in polystyrene, flat-bottom 96-well plates and were allowed 24 hours to adhere at 37°C in a 5% CO2 incubator. Cells were then treated with 5-fluorouracil (48.8nM to 200uM), F10 (73.2pM to 300nM), or corresponding vehicle controls. 200uM exogenous uridine, thymidine, or PBS was then added to the respective wells and gently mixed as to not disturb cellular growth. After 10 days, cellular viability was determined through resazurin assays as performed in the earlier drug screens.

### Generation of TP53-KO minor and major MCF7 cell lines via CRISPR-Cas9

TP53-KO minor and major MCF7 cells were generated using the previously published TCGI System (Zhao et al., 2019). In brief, a gBlock containing sgRNA4: 5’-GGACGATATTGAACAATGG-3’ and sgRNA5: 5’-GGGCAGCTACGGTTTCCGTC-3’ was amplified by PCR using the following primers: TCGI_Amplifying_F: 5’-AGGCACTTGCTCGTACGACG-3’ and TCGI_Amplifying_R: 5’-ATGTGGGCCCGGCACCTTAA-3’. The PCR product was then gel purified with NEB Monarch^®^ DNA Gel Extraction Kit (NEB, Cat No. T1020). Alongside pLentiCRISPRv2 (Addgene, Cat. No. 52961), the construct was digested with BsmBI (NEB, Cat. No. R0580) at 55°C for 1 hour, after which the enzyme was heat-inactivated at 80°C for 20 minutes (Sanjana et al., 2014). The digested constructs were then purified with Ampure XP Beads and then ligated with the Quick Ligation Kit (NEB, Cat. No. M2200S).

Lentiviral packaging mix was produced from 2.5μg of TP53-KO pLentiCRISPRv2 construct or vector backbone, 0.625ug pMD2.G (Addgene, Cat. No. 12259), and 1.875ug psPAX2 (Addgene, Cat. No. 12260). The packaging mixes were transfected into HEK293T cells with Lipofectamine3000 to produce TP53-KO lentivirus and EV (Empty Vector) control lentivirus. Both lentiviruses were used to transduce MCF7 cells in cellular media containing 10ug/mL polybrene (Sigma-Aldrich, Cat. No. TR-1003-G). After 24 hours of transduction, old media was replaced with polybrene-free media for 48 hours before the cells underwent puromycin (ThermoFisher, Cat. No. A1113802) selection at 2μg/mL for 7 days to obtain the TP53-KO minor cell line. To produce the TP53-KO major cell line, the puromycin-selected MCF7 cells were cultured in 20uM Nutlin-3a for 1 week.

### Matrigel drug screening in TP53-KO minor and major MCF7 cells

MCF7 TP53-EV and TP53-KO minor/major cells were first resuspended in a slurry consisting of 3 parts cell media and 4 parts matrigel to obtain a density of 50 cells/μL slurry. Both cell slurries were then plated at a density of 1000 cells/well in polystyrene, flat-bottom 96-well plates and allowed to solidify at 37°C in a 5% CO2 incubator. Once solidified, 200μL of cell media was added to cover the cells embedded in matrigel, and the cells were returned to the incubator. After 12 hours, normal cell media was replaced with media containing drugs or corresponding vehicle controls at concentrations ranging from 73.24pM to 300μM. Drug media was replaced every 3 days for 30 days, after which cellular viability was quantified using the Cell-TiterGlo 2.0 Viability Assay (Promega, G9241).

### Sequencing of TP53 Genomic Locus in TP53-KO minor MCF7 cells

Genomic DNA was isolated from all TP53-KO minor cells treated in F10 matrigel assays using the Agencourt DNAdvance Genomic DNA Isolation Kit (Beckman Coulter, Cat. No. A48705), and the TP53 locus was amplified using the following primers: TP53_SeqF: 5’-CCTGGTCCTCTGACTGCTCT-3’ and TP53_SeqR: 5’-GCCAAAGGGTGAAGAGGAAT-3’. Nanopore sequencing was then performed on the PCR products with the Ligation Sequencing Kit (ONT, Cat. No. SQK-LSK109) and a MinION Flow Cell (ONT, Cat. No. R9.4.1). Reads were mapped to reference, and CRISPResso2 was utilized to identify the 7 most frequent indels present in vehicle control treated TP53-KO minor cells (Clement et al., 2019). Counts for these 7 specific indels were then obtained in F10-treated wells using the “ugrep” command on a Mac Terminal. For plotting, indels fractions were calculated and subsequently normalized to vehicle control measurements.

## Results

### Generation and characterization of pooled TP53-KO MCF7 cells

TP53-mutant luminal A (HR+/PR+/HER2-) invasive ductal carcinomas have strongly correlated with decreased patient survival in the metastatic setting (Fig. 1A), and for this reason we selected the representative, TP53-WT MCF7 breast adenocarcinoma cell line for our studies (Cerami et al., 2012; Gao et al., 2013; Nguyen et al., 2022; Dai et al., 2017). Given that approximately 91% of documented TP53 mutations in the TCGA Pan-Cancer Cohort resulted in loss of WT p53 function, we chose the CRISPR/Cas9 homozygous deletion system to model the functional effects of TP53 mutation in cancer and screen for TP53-associated drug sensitivity in MCF7 cells (Donehower et al., 2019). We first generated two unique pooled TP53-KO MCF7 derivatives using CRISPR/Cas9 genomic engineering: TP53KO 3.4 and TP53KO 5.6 (Fig 1B). Pool 3.4 was produced from sgRNAs targeting Exon 4 and Exon 10, and pool 5.6 was produced from sgRNAs targeting Exon 4. After co-transfection of a gBlock encoding both sgRNAs and a Cas9-encoding plasmid, p53-null cells were selected for by treatment with 10uM Nutlin-3a, a competitive inhibitor of the p53-MDM2 interaction and a potent activator of p53-induced apoptosis. After selection, both TP53-KO pools were resistant to Nutlin-3a over a 10-day time-course treatment, indicating phenotypic behavior consistent with a TP53-KO profile (Fig. 1C).

**Figure 1.**
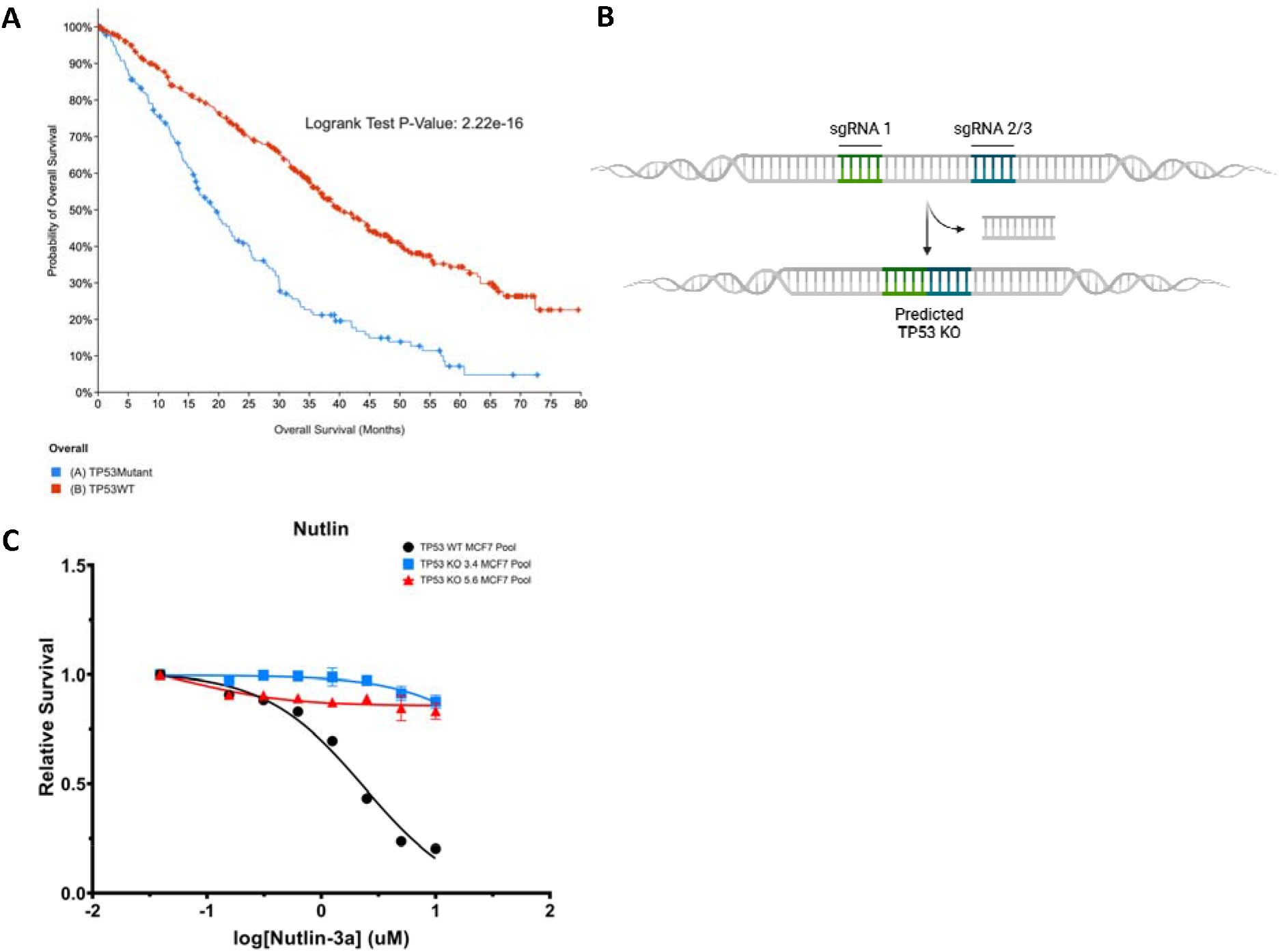
Generation of pooled TP53-KO MCF7 cells via CRISPR-Cas9 genomic engineering. **A)**Overall survival curves of metastatic HR+/HER2-ductal breast cancer patients stratified by TP53-status in the MSK-MET cohort. Data was obtained from www.cbioportal.org. **B)** Schematic depicting the relative sgRNA target locations within the TP53 locus designed to produce gene deletions. **C)** Functional assay for phenotypic TP53-KO behavior using Nutlin-3a, a competitive inhibitor of p53-MDM2 interaction

### Loss of p53 promotes sensitivity to multiple chemotherapeutic agents in MCF7 cells

To determine whether TP53-status affected chemosensitivity, we screened the NCI Approved Oncology Drug Set IV (Supplementary Table S1) against both the TP53-KO pools and the TP53-WT MCF7 cells (Fig. 2A). Dose-response measurements were performed in triplicate after 10 days of drug treatment using a fluorescent resazurin viability assay. While most drugs displayed no TP53-dependent response to treatment, we found 6 drugs to elicit significant differential responses between TP53-KO and TP53-WT cells (Fig. 2B). Nutlin and oxaliplatin were both less effective on p53-null cells, consistent with documented clinical correlations between TP53 mutation and resistance to DNA damaging chemotherapy (Liu et al., 2019). Surprisingly, we also found 4 antimetabolites/DNA synthesis inhibitors (5-FU, gemcitabine, cytarabine, clofarabine) that preferentially targeted TP53-KO cells over TP53-WT cells.

**Figure 2.**
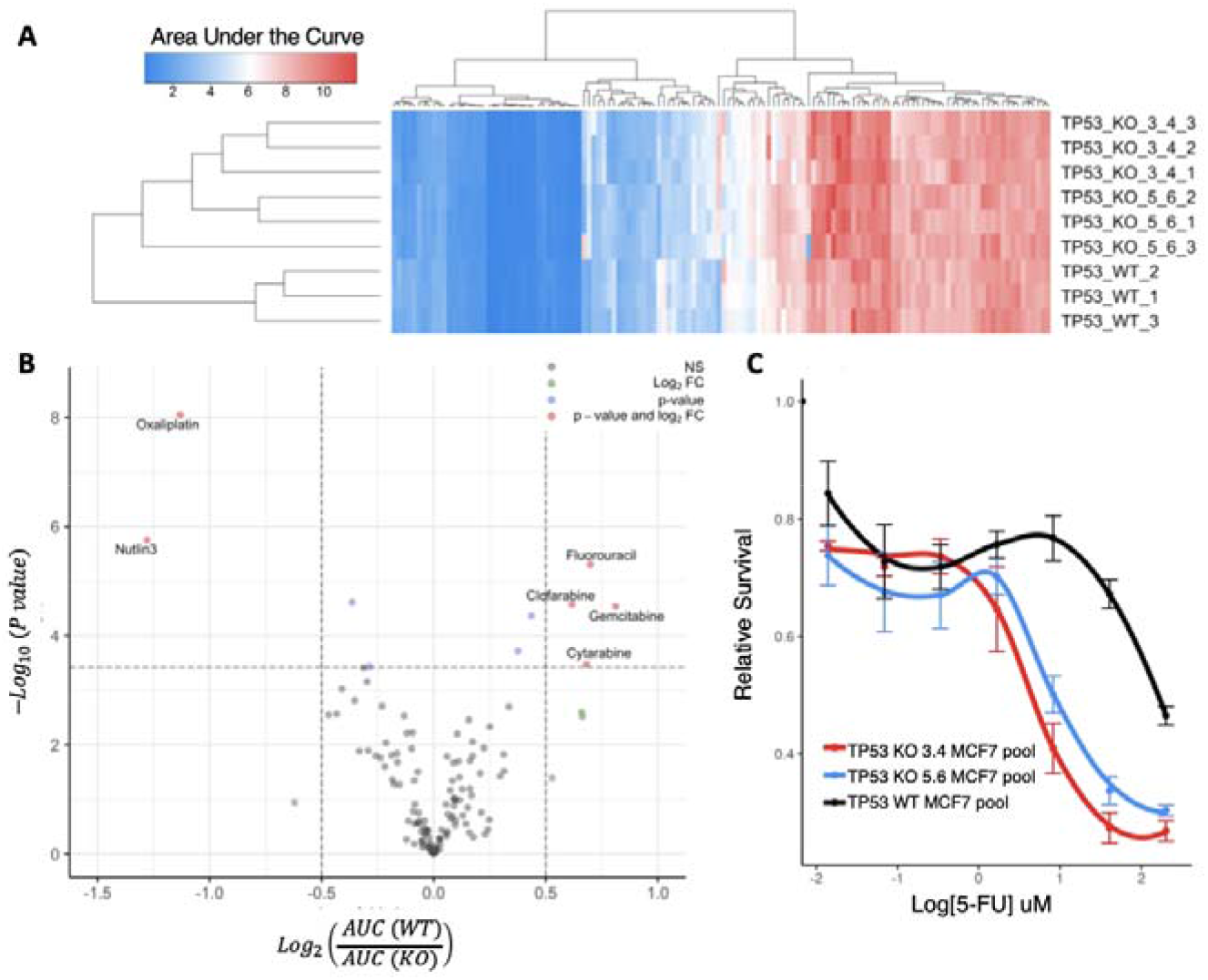
Systematic screen of 129 FDA-approved oncology drugs in pooled TP53-KO and TP53-WT MCF7 cells. **(A)** The heat map of AUC for all drugs screened in the study. Each drug represents a column and each row represents a cell line (in triplicate). The color of each cell corresponds to the AUC – a surrogate measure of drug resistance. **(B)** A volcano plot representation of Student t test results depicting the magnitude (log2 of ratio between AUC of TP53-WT and of TP53-KO, x-axis) and significance (p-value, y-axis) of TP53-associated drug sensitivity. Each drug is represented by a dot, and significant drugs (p < 0.01, and fold change (FC) > 1.25) are colored red. **(C)** Viability curve of TP53-WT and KO pools treated with 5-FU in drug screen.

### F10 offers increased cytotoxicity in p53-null MCF7 cells via a DNA-directed mechanism of action

Because 5-FU displayed the most significant preferential toxicity towards TP53-KO MCF7 cells in the drug screen (Fig. 2B-C), we hypothesized that second-generation fluoropyrimidines might induce greater cytotoxicity in p53-null cells than 5-FU. To investigate this, we first revalidated 5-FU’s preferential toxicity observed in the drug screen in single-cell clones isolated from both TP53-KO pools and TP53-WT MCF7 cells to ensure our initial results had not been biased by aberrant clones in the pools (Fig. 3A). Upon confirmation, we treated the clones with F10, a novel polymeric fluoropyrimidine. To compare the chemosensitivity of F10 to 5-FU, we calculated the fold-change in AUC difference for TP53-WT and TP53-KO clones treated with both drugs. After a 10-day time-course assay, we observed a 6.11 and 3.00-fold increase in TP53-KO sensitivity for F10-treated clones compared to 5-FU treated clones (Fig. 3A, p < 0.00001).

**Figure 3.**
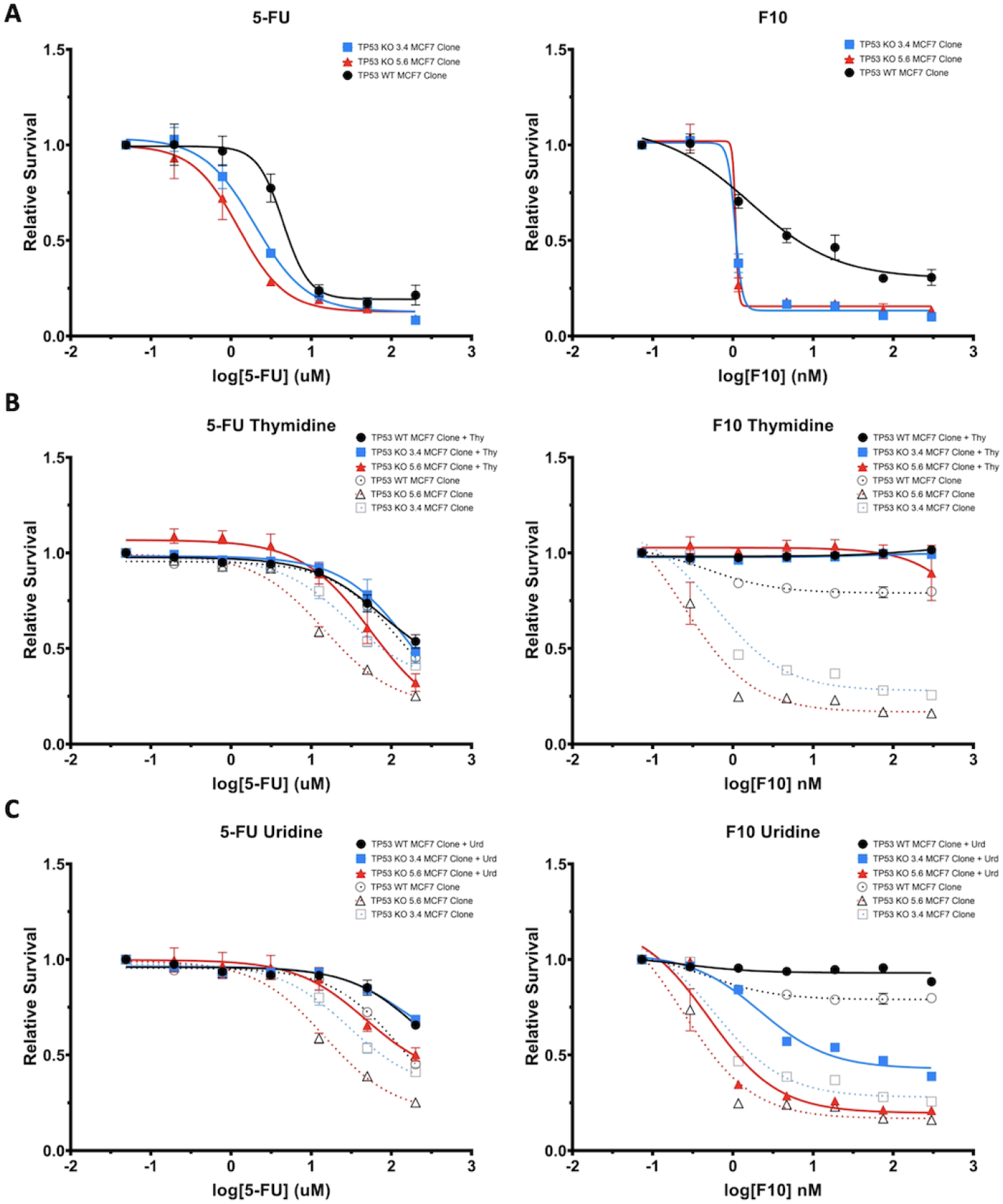
DNA-directed mechanism of F10 greatly impairs TP53-KO cells. **(A)** Viability curves of 5-FU and F10-treated clonal TP53-KO and TP53-WT MCF7 cells. **(B)** Viability curves of thymidine and **(C)** uridine rescue assays on 5-FU and F-10 treated clonal TP53-KO and TP53-WT MCF7 cells.

To further investigate the mechanism behind F10’s increased synthetic lethality with TP53-KO in MCF7 cells, we performed rescue experiments with exogenous uridine and thymidine on 5-FU and F10-treated clones (Fig. 3B). Here, we observed that 5-FU’s preferential killing of TP53-KO cells could only partially be attributed to its incorporation into DNA or RNA, whereas F10’s preferential killing of TP53-KO cells could be completely abrogated by the addition of exogenous thymidine, with only minimal rescue observed by exogenous uridine. These results suggest DNA incorporation is central to F10’s cytotoxic mechanism.

### F10 effects on TP53-KO cells are persistent in heterogeneous settings reflective of the tumor microenvironment

Given that TP53 mutation has been suggested to be an initiating event present at all stages of luminal A tumorigenesis, we were interested in determining whether F10 could selectively eliminate TP53-KO cells from both subclonal (TP53-KO minor) and clonal (TP53-KO major) cellular populations (Bertheau et al., 2013). To investigate this, we adapted the tRNA-CRISPR for Genetic Interactions (TCGI) system in MCF7 cells to produce both populations by selecting for the TP53-KO phenotype with Nutlin-3a and/or integration of the lentiviral vector with puromycin (Fig. 4A). In a 30-day matrigel F10 time-course assay, we observed a lower IC_50_ for the TP53-KO major population than the TP53-KO minor population, which is consistent with the expected proportion of TP53-KO cells in each pool (Fig. 4B). To rule out the possibility of the observed killing being related to remaining TP53-WT cells in the pooled populations, we performed nanopore sequencing on genomic DNA extracted from TP53-KO minor samples at the end of the time-course assay. We observed significant disappearance of the 7 most frequent TP53-KO indels, as identified with CRISPRessov2, with addition of the lowest concentration of drug (0.286pM F10).

**Figure 4.**
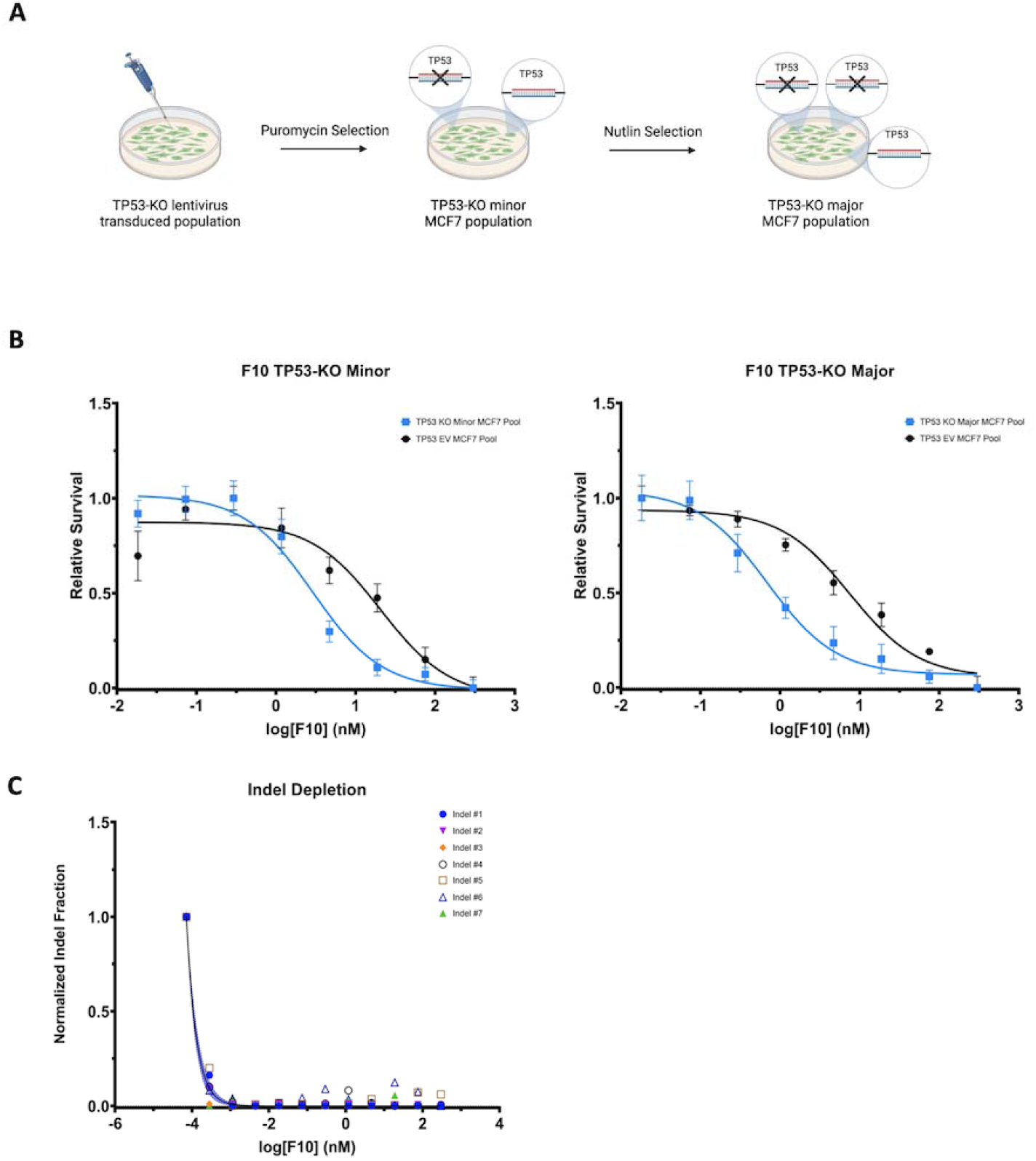
F10 can selectively target TP53-KO cells in heterogeneous cell populations. **(A)** Schematic depicting workflow to produce TP53-KO minor and major MCF7 cell populations. **(B)** Viability curves of TP53-KO minor and major and TP53-EV MCF7 cells treated with F10 in matrigel assays. **(C)** Indel fraction after F10 treatment of the 7 most frequent indels produced by CRISPR/Cas9 genomic engineering in the TP53-minor MCF7 cell population. Indel fractions are normalized to vehicle control.

## Discussion

Given that TP53 is the most frequently mutated gene in human cancers, it has always been considered an exceptional target for therapeutic development. Many projects have aimed to produce gene therapies, identify synthetic lethal interactions, or design immunotherapies targeting mutant p53 with the hope of benefiting patients across all cancer types (Zhang et al., 2018; Wang & Simon, 2013; Hsiue et al., 2021). Our previous studies investigated the p53-dependency of drug sensitivity in human embryonic stem cells and colorectal cancer cells, and here, we aimed to extend our analyses to breast cancer models (Liu et al., 2019). We generated TP53-KO MCF7 breast cancer cells, investigated their sensitivity to 129 FDA-approved chemotherapeutics, and identified 5-fluorouracil to confer the highest degree of selective TP53-KO killing. We then examined the effects of a novel polymeric fluoropyrimidine, F10, on the p53-null cells and found F10 to be significantly more potent, via a DNA-directed mechanism of action.

Resistance to chemotherapy can be a major hurdle in treating cancer patients, and several pre-clinical and clinical studies have shown TP53 mutations can be predictive markers of chemoresistance (Ferreira et al., 1999). For example, in ovarian cancer, p53 inactivation is associated with resistance to traditional platinum-based therapies, and in breast cancer, p53 inactivation has been linked with anthracycline resistance (Reles et al., 2001; Aas et al., 1996). Our drug screen surprisingly found p53-null breast cancer cells to be chemosensitive to several antimetabolites, the most significant of which was 5-fluorouracil, a drug routinely used in the clinical management of breast cancer patients.

Despite its widespread use alongside cyclophosphamide and methotrexate, 5-FU treatment may not always synergize with TP53-mutation in breast cancer (Waks & Winer, 2019; Geisler et al., 2003). In particular, Geisler et al. have reported that TP53-mutation could actually serve as a predictor of chemoresistance to a 5-FU/mitomycin regimen in a cohort of locally advanced breast cancer patients. Although they were assessing a combinatorial regimen, and the accuracy of TP53-mutant identification has markedly increased since their study, their work still illustrates the necessity of testing novel therapeutics for synergy with TP53-KO. Given the polymeric structure of the novel fluoropyrimidine, F10, we hypothesized that the p53-null cells would be even more sensitive to F10 than 5-FU. Our results clearly indicate F10’s dramatic increase in selective TP53-KO killing relative to 5-FU across multiple genetic TP53-KO signatures and in settings more reflective of the tumor microenvironment.

This increased potency of F10 has been documented in other tumor types, including AML, ALL, GBM, colorectal, and prostate cancer, and is primarily linked to the increased conversion of F10 to FdUMP, relative to 5-FU (Pardee et al., 2012, 2014; Gmeiner, Lema-Tome, et al., 2014; Gmeiner et al., 2021; Gmeiner, Willingham, et al., 2014; Gmeiner et al., 2016). FdUMP is released from F10 via the activity of 3’-O-exonucleases, thus circumventing the need for metabolic activation that is required for the conversion of 5-FU to FdUMP (Liao et al., 2005). These increased levels of FdUMP allow for greater inhibition of Thimidylate Synthase (TS) and subsequent elevated incorporation of dUTP and FdUTP in DNA, resulting in “thymineless death” (Longley et al., 2003). Our Urd/Thy rescue experiments demonstrate that F10’s selective killing of TP53-KO cells is mediated by a DNA-directed mechanism of action. The lack of significant RNA-dependent effects suggests that F10 may help curb systemic gastrointestinal toxicities associated with 5FU’s RNA-directed mechanisms (Pritchard et al., 1997).

In conclusion, we generated TP53-KO breast cancer cells, screened them with 129 FDA-approved oncology drugs, and identified the p53-null cells to be highly sensitized to 5-FU. This chemosensitivity was dramatically increased by treatment with the novel polymeric fluoropyrimidine F10, via a DNA-directed mechanism of action. Together, our findings indicate that F10 may offer an improved therapeutic window for TP53-mutant breast cancer patients.

## Disclosures of Potential Conflicts of Interest

PB reports receiving a commercial research grant from Janssen Research & Development. No potential conflicts of interest were disclosed by other authors.

## Notes

### Competing Interest Statement

The authors have declared no competing interest.

